# Long-Range Electrostatic Interactions Significantly Modulate the Affinity of Dynein for Microtubules

**DOI:** 10.1101/2021.11.24.469892

**Authors:** Ashok Pabbathi, Lawrence Coleman, Subash Godar, Apurba Paul, Aman Garlapati, Matheu Spencer, Jared Eller, Joshua Alper

**Affiliations:** Department of Physics and Astronomy, Clemson University, Clemson, SC 29634; Eukaryotic Pathogen Innovations Center, Clemson, SC 29634; School of Mathematical and Statistical Sciences, Clemson, SC 29634; Department of Genetics and Biochemistry, Clemson, SC 29634; Department of Biological Sciences, Clemson, SC 29634

**Author notes:** Department of Industrial Chemistry, Mizoram University, Aizawl, Mizoram, India 796004 (Ashok Pabbathi). Department of Physics, University of Maryland Baltimore County, Baltimore, MD 21250. Department of Electrical Engineering, University of Notre Dame, Notre Dame, IN 46556. Department of Biochemistry, University of Texas at Austin, Austin, TX 78712.

## Abstract

The dynein family of microtubule minus-end directed motor proteins drives diverse functions in eukaryotic cells, including cell division, intracellular transport, and flagellar beating. Motor protein processivity, which characterizes how far a motor walks before detaching from its filament, depends on the interaction between its microtubule-binding domain (MTBD) and the microtubule. Dynein’s MTBD switches between high- and low-binding affinity states as it steps. Significant structural and functional data show that specific salt bridges within the MTBD and between the MTBD and the microtubule govern these affinity state shifts. However, recent computational work suggests that non-specific, long-range electrostatic interactions between the MTBD and the microtubule may also play a significant role in the processivity of dynein. To investigate this hypothesis, we mutated negatively charged amino acids remote from the dynein MTBD-microtubule-binding interface to neutral residues and measured the binding affinity using microscale thermophoresis and optical tweezers. We found a significant increase in the binding affinity of the mutated MTBDs for microtubules. Furthermore, we found that charge screening by free ions in solution differentially affected the binding and unbinding rates of MTBDs to microtubules. Together, these results demonstrate a significant role for long-range electrostatic interactions in regulating dynein-microtubule affinity. Moreover, these results provide insight into the principles that potentially underlie the biophysical differences between molecular motors with various processivities and protein-protein interactions more generally.

**Statement of Significance:** The dynein family of motor proteins drives the motility of multiple cellular functions by walking toward the minus end of microtubules. The biophysical mechanisms of dynein rely on its ability to change affinity for the microtubule as it steps. Specific short-range electrostatic interactions acting at the microtubule-binding domain (MTBD)-microtubule interface are known to govern binding affinity. This study shows that non-specific longer-range electrostatic interactions due to charged amino acids remote from the binding interface also contribute significantly to the binding affinity mechanisms. Our results suggest that subtle differences in the electrostatic charge distribution within the MTBD significantly affect the molecular biophysical motility mechanisms in the dynein family of motors.

## Introduction

Cytoplasmic dynein (hereafter called *dynein* unless otherwise specified) is a microtubule minus-end directed motor protein that drives diverse functions in eukaryotic cells, including retrograde intracellular transport (1), mitotic spindle assembly (2), chromosome segregation (3), nucleus positioning (4, 5), and cytoskeletal network organization (6). Dynein dysfunction is associated with several diseases, in particular, neurodegenerative diseases, including Alzheimer’s, Parkinson’s, and Huntington’s disease (7–15). Cytoplasmic dynein is a 1.4 MDa protein complex with two copies of a 530 kDa heavy chain and associated regulatory light, intermediate light, and intermediate chains (16). The dynein heavy chain consists of a dimerizing tail domain, a linker, and a microtubule-binding domain (MTBD) separated from a ring of six AAA^+^ domains by a 15 nm coiled-coil stalk (Fig. 1 *A*) (16, 17). Communication between the ATPase active site in AAA1 and MTBD through the coiled-coil stalk modulates the binding affinity of dynein towards microtubules (18, 19).

**Figure 1.**
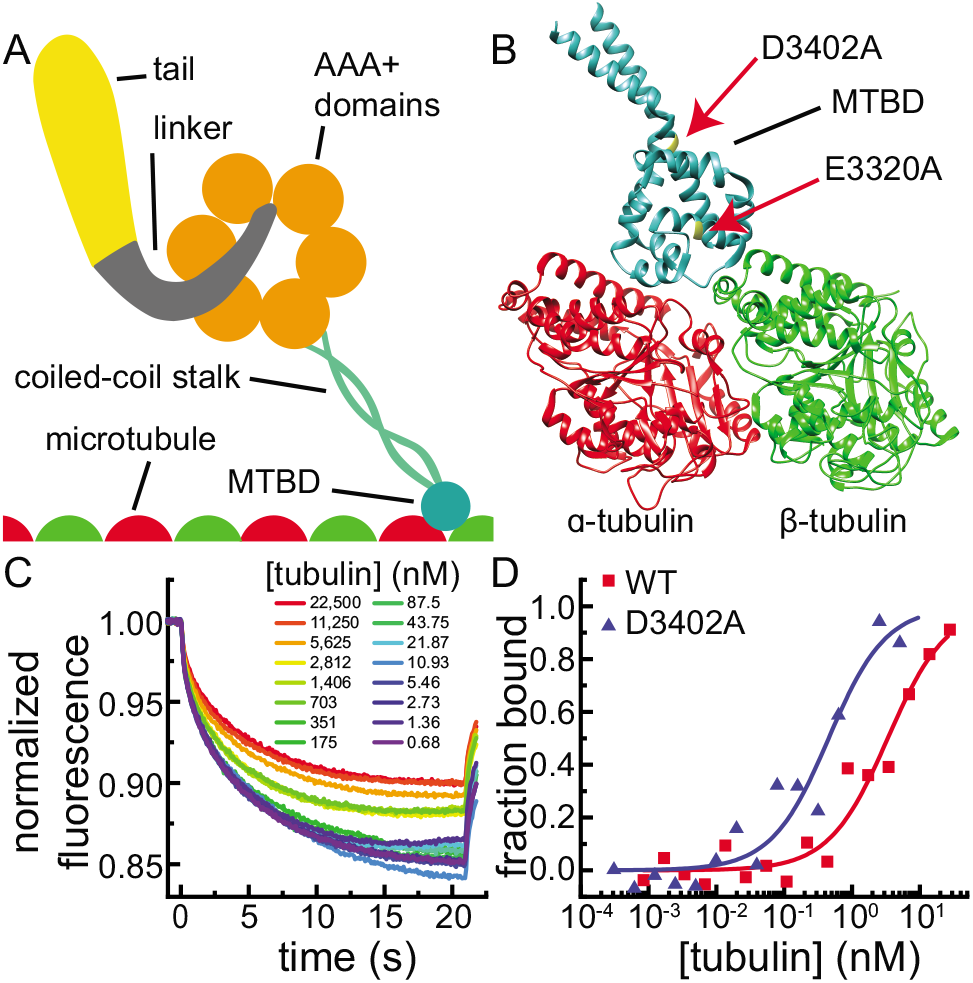
D3402A mutant MTBDs have a higher binding affinity for microtubules than wild-type MTBDs. (*A*) Schematic representation of a cytoplasmic dynein heavy chain bound to a microtubule with its principal domains indicated. (*B*) Ribbon diagram representation of MTBD (*teal*) shown bound to a tubulin dimer (high-binding affinity dynein microtubule-binding domain - tubulin complex cryo-EM, PDB ID: 3J1T (24)). The locations of E3320A and D3402A mutations, which were chosen to be remote from the binding interface to investigate the role of long-range electrostatic interactions, are indicated (*highlight* with *red arrows*). (*C*) MST traces for WT MTBDs in 0 mM KCl. Each MST trace corresponds to a concentration ranging from 0.68 nM (*violet*) to 22,500 nM (*red*) of tubulin dimers polymerized into the microtubules. The fluorescence of each trace is normalized by the fluorescence when temperature gradient inducing laser irradiation starts (at time = 0). (D) MST-derived Hill–Langmuir binding curves for both WT (*red line*) and D3402A (*blue line*) MTBDs in 0 mM KCl fit to the fraction bound data (*red circles* and *blue triangles*, respectively). The *K*_d_s were 4.48 ± 0.29 μM for WT and 0.39 ± 0.06 μM (fit parameter ± standard error of the fit) for WT and D3402A, respectively. The fluorescently-labeled MTBD concentration was 50 nM and the concentration of microtubules varied from 0 to 23 μM in both cases.

Processivity, which is a measure of how far a motor protein moves along its associated filament before detaching (20), is an essential biophysical property affecting motor proteins’ cellular functionality. Dynein processivity depends on the details of the affinity of the MTBD for the microtubule (21); it must switch between high- and low-binding affinity states as a function of ATP hydrolysis state in a coordinated way to walk along a microtubule (21, 22). The affinity of dynein for microtubules switches between high- and low-binding affinity states due to conformational changes in the MTBD, which results from changing registrations of the coiled-coil stalk domain helices (19, 23–26) in response to the ATPase cycle state in AAA1 (19, 22, 27, 28). It is thought that dynein switches to its high-binding affinity state when it is unbound from the microtubule, causing it to bind, and that it switches to its low binding affinity state when it is bound to the microtubule, causing it to unbind and step forward (29, 30).

Electrostatic interactions dominate the dynein-microtubule binding affinity mechanisms. Charged-to-neutral and charge-flipping amino acid substitutions in the MTBD's microtubule-binding interface altered the microtubule-binding affinity (22, 29, 31), presumably due to changes in surface charge complementarity (32, 33). Additionally, multiple dynamic salt-bridges within the MTBD and between the MTBD and the microtubule regulate the MTBD-microtubule-binding affinity (24). Together, these results indicate the importance of both specific and non-specific electrostatic interactions in the biophysical mechanisms of dynein.

Previously, our computational work on the dynein MTBD-microtubule-binding interface suggested that non-specific, long-range electrostatic interactions contribute significantly to the MTBD-microtubule-binding affinity (34). When we computationally mutated multiple charged amino acids that are remote from the binding interface to neutral ones, we found that the changes in the electrostatic component of binding energies are similar in magnitude to changes associated with mutations to the previously identified specific dynamic salt bridges (34). Moreover, our computational investigation of interactions between tubulin's highly-charged C-terminal tails (CTTs or E-hooks) and the MTBD further indicated the importance of long-range, non-specific electrostatic interactions in guiding dynein-microtubule binding (35). Therefore, we hypothesize that long-range electrostatic interactions, i.e., those remote from the binding interface, between the MTBD and the microtubule contribute significantly to the motility mechanisms of dynein.

To investigate the effect of long-range electrostatic interactions on the binding of dynein to microtubules, we characterized the binding affinity of wild-type and long-range electrostatic interaction-altering mutant MTBDs (Fig. 1 *B*) for microtubules in the high-binding affinity (22) state based on our computational work (34). We measured the equilibrium binding affinity using microscale thermophoresis (MST) and the force-dependent unbinding rate using optical tweezers in varying concentrations of added salt. We found that mutating a negatively charged amino acid remote from the binding interface to a neutral one enhanced the binding affinity and reduced the force-dependent unbinding rate of the MTBD in an ionic strength-dependent manner. Together, our results suggest a significant contribution of long-range electrostatic interactions to regulating dynein’s processivity.

## Materials and Methods

### Tubulin purification and labeling

Tubulin was purified from porcine brains using a phosphocellulose column (PC-tubulin, gift from Jonathon Howard, Yale University) (36). We cycled (polymerized and depolymerized) and rhodamine-labeled the PC-tubulin using standard protocols (37). In brief, we incubated a 10-fold excess of rhodamine dye (5(6)-TAMRA, SE, Biotium, Fremont, CA) with a microtubule solution in HEPES labeling buffer (0.1 M NaHEPES, pH 8.6, 4 mM MgCl_2_, 1 mM EGTA, 40% (v/v) glycerol) for 40 min at 37°C. We then depolymerized the labeled microtubules at 4°C and pelleted any remaining microtubules. We cycled the labeled tubulin again to ensure all the labeled tubulin was polymerization competent. We measured the degree of labeling from the concentration of tubulin and the TAMRA dye using a UV-Vis spectrophotometer (NanoDrop One, Invitrogen, Waltham, MA) with an extinction coefficient for tubulin at 280 nm of 115,000 M^-1^cm^-1^, a correction factor of 0.3, and an extinction coefficient for the TAMRA dye at 555 nm of 65,000 M^-1^cm^-1^. We found that the degree of labeling was 0.8 rhodamine molecules per tubulin dimer.

### Microtubule preparation

We prepared taxol-stabilized, unlabeled microtubules using standard methods, with slight modifications (38). Briefly, we added unlabeled tubulin (32 μM) to polymerization buffer (4% DMSO, 4 mM MgCl_2_, 1 mM GTP, and BRB80 (80 mM PIPES, 1 mM EGTA, 1 mM MgCl_2_ at pH 6.9)) for 30 min at 37°C to polymerize microtubules. We diluted the microtubules in BRB80T (BRB80 with 20 μM Taxol (paclitaxel, J62734, Alfa Aesar, Tewksbury, MA)) and pelleted them in an air-driven centrifuge at 30 psi (Airfuge, Beckman Coulter, Inc., Indianapolis, IN). We resuspended the pellet in BRB80, took a small fraction of the microtubules from the solution for further analysis, and added 20 μM Taxol to the rest of the solution. We depolymerized the small fraction of microtubules at 4°C and measured the concentration of tubulin at 280 nm using the NanoDrop.

We prepared polarity-marked microtubules as described previously with some modifications (39). Briefly, we prepared GMPCPP-stabilized bright microtubules seeds by incubating 2 μM rhodamine-labeled tubulin in 1 mM GMPCPP (guanosine-5'-[(α,β)-methyleno]triphosphate, sodium salt, GMPCPP, NU-405, Jena Bioscience, Jena, Germany), 1 mM MgCl_2_, and BRB80 for 30 min at 37 °C. We pelleted the microtubule seeds at 30 psi in the airfuge, resuspended them in 50 μL of BRB80, and sheared using a high-gauge needle to make short, highly-labeled seeds. We made dim extensions to the bright seeds by incubating the seeds with 2.2 μM tubulin (5:1, unlabeled:labeled tubulin mixture), 1 mM GTP, 1 mM MgCl_2_, and BRB80 for 2 hours at 37 °C. Afterward, we added 350 μL of BRB80, pelleted microtubules were at 30 psi in the airfuge, resuspended in 100 μL BRB80T, and used them within a week. We identified the plus-ends of the polarity-marked microtubules as having longer dim extensions from the bright nucleating seeds (39).

### MTBD expression and purification

The SRS-MTBD (22/19), AB (monomer), wt plasmid, which encodes a chimeric seryl-tRNA synthetase (SRS) – mouse cytoplasmic dynein MTBD fusion protein (22), was a gift from Ron Vale (Addgene plasmid # 22380; http://n2t.net/addgene:22380; RRID:Addgene_22380). The SRS domain located at the distal end of the coiled-coil stalk serves to stabilize and lock the registry stalk (22). This 22/19 construct has 22 and 19 amino acids between the registration-locking SRS domain and the conserved proline residues in each strand of the coiled-coil stalk (22). The 22/19 construct represents the high-binding affinity stalk registration (22). We mutated aspartic acid 3420 to alanine (D3420A) and glutamic acid 3320 to alanine (E3320A) using a Q5 site-directed mutagenesis kit (E0552S, New England Biolabs, Ipswich, MA, Table S2 in the Supporting Material for primer sequences). These mutations changed negatively charged amino acids remote (2.6 nm and 1.2 nm, respectively, measured in PDB ID: 3J1T (24)) from the binding interface (Fig. 1 *B*) to neutrally charged residues, as suggested in our previous computational study (34).

We expressed the MTBD constructs in *E. coli* by overnight induction with 0.5 mM IPTG at 18 °C. We harvested the cells by centrifugation (4000 ×g for 25 min), resuspended them in binding buffer (20 mM sodium phosphate, 500 mM NaCl, and 40 mM imidazole, pH 7.4), and lysed using an ultrasonic tip sonicator. We clarified the lysate by centrifugation (4000 × g for 35 min) and purified the MTBD constructs from the lysates using a HisTrap High Performance nickel Sepharose column (17-5247-01, Cytiva (formerly GE Healthcare Life Sciences), Marlborough, MA), per the manufacturer’s instructions. Briefly, we loaded the sample onto the column, washed it with 5 column volumes, and eluted the 6×His-tagged MTBDs with an elution buffer (20 mM sodium phosphate, 500 mM NaCl, and 500 mM imidazole, pH 7.4) in 1 column volume fractions. We pooled fractions containing the purified protein, desalted the pooled fractions using PD-10 desalting columns packed with Sephadex G-25 resin (17085101, Cytiva (formerly GE Healthcare Life Sciences), Marlborough, MA), and stored aliquots in PBS with 10% glycerol at -80℃. Densitometry of Coomassie blue-stained SDS-PAGE gel electrophoresis showed approximately 90% purity of each MTBD construct used (Fig. S1 in the Supporting Material).

To test the long-range electrostatic interaction hypothesis, the charge-altering mutations remote from the binding interface must not change other significant contributing factors to binding affinity, like the structure. We assessed the structural stability of the MTBD constructs to mutations in these charged residues using circular dichroism (Fig. S2 in the Supporting Material) and computational methods (DynaMut (40) and Site Directed Mutator (SDM) (41), Fig. S3 in the Supporting Material). We found that the D3402A mutation did not destabilize the MTBD but that the E3320A mutation caused significant structural changes (see Supporting Material for details). Therefore, we proceeded to make binding affinity and dissociation measurements with the D3402A construct only.

### Microscale thermophoresis binding affinity measurements

Microscale thermophoresis (MST) measures the dissociation constant, *K*_d_, of a biomolecular interaction based on changes in thermophoretic mobility, which is the directed movement of molecules in a microscopic temperature gradient when one fluorescently-labeled molecule binds to an unlabeled binding partner (42). We used MST because it requires significantly less sample than alternative techniques like isothermal calorimetry (42).

We measured the binding affinity of MTBDs (fluorescently labeled target) to microtubules (unlabeled ligand) using a Monolith NT.115 MST system (NanoTemper Technologies, GmbH, Munich, Germany). We labeled the MTBDs with a 2nd Generation RED-tris-NTA His-Tag Labeling Kit (MO-L018, NanoTemper Technologies, GmbH, Munich, Germany) per the manufacturer’s instructions with slight modifications (43). In brief, we mixed equal volumes of purified MTBD protein (200 nM) and RED-tris-NTA dye (100 nM) and incubated overnight at 4°C. Due to the excess of MTBD and the high affinity of the dye for the MTBD’s His-Tag, these labeling conditions ensured that the dye molecules bound to the MTBDs in 1:1 stoichiometry (43). Then, we removed protein and dye aggregates from the sample by centrifugation (18,000 × g) at 4°C for 10 minutes. We diluted the supernatant, which consisted of labeled MTBD, to 50 nM and proceeded with MST.

To perform MST, we prepared a 16-step two-fold serial dilution series of Taxol-stabilized, unlabeled microtubules. The series started with approximately 23 μM and ended with approximately 0.6 nM tubulin dimers polymerized in the microtubules. We incubated each sample from the serial dilution, as well as a no microtubule control, with 50 nM of RED-tris-NTA labeled MTBD in BRB80T with 0.2% IGEPAL CA-630 (J61055, Alfa Aesar, Tewksbury, MA). To further probe the effect of long-range electrostatic interactions on the dissociation constant, we performed MST in samples with 0 mM, 100 mM, and 300 mM of KCl added. These salt concentrations decrease the Debye length, i.e., the characteristic length scale by which the electrostatic effects decay in the solvent, from 0.8 nm to 0.6 nm and 0.4 nm, respectively, per

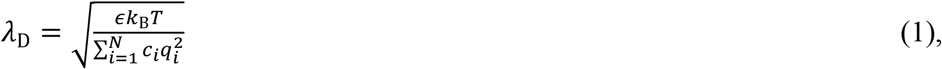

where *ϵ* is the dielectric constant of the solvent (80.4 for water), *k*_B_*T* is the scale factor for molecular energy (Boltzmann’s constant times absolute temperature = 4.11 × 10^-21^ J), *c*_*i*_ is the concentration of charged species, *i*, *q*_*i*_ is the charge carried by species, *i*, and *N* is the total number of mobile charged species in solution. However, the additional KCl is not a sufficient concentration to cause a significant Hofmeister (44) and Kirkwood “salting out” effect (45, 46).

We transferred the samples to glass capillaries tubes (MO-K022, NanoTemper Technologies, Munich, Germany). We acquired MST traces, which track fluorescence intensity changes due to thermophoresis in a confocal laser-induced temperature gradient within the capillaries (42), at 40% excitation power of Nano-RED LED. We fit the traces to a linearized model of MST, found the fraction of bound MTBD, and calculated the *K*_d_ of the fluorescently labeled target MTBDs for the unlabeled ligand microtubules using the Hill–Langmuir equation (47), as provided in the MO.Affinity analysis software (NanoTemper Technologies, Munich, Germany).

### Conjugation of MTBD to beads for force-dependent dissociation optical tweezer assays

We used a biotinylated anti-His Tag antibody-streptavidin method to conjugate the MTBDs to polystyrene beads for the force-dependent dissociation optical tweezer assays (48). Briefly, we washed 1% (w/v) 1.04 μm diameter, streptavidin-coated polystyrene microspheres (CP01004, Bangs Laboratories, Inc., Fishers, IN) three times with centrifugation (11,000 × g, 3 min) and resuspension in PBS with 0.1% Tween 20 (V0777, AMRESCO LLC., Solon, OH). We incubated the washed beads with 0.25 mg/mL mouse monoclonal biotinylated anti-His Tag monoclonal antibody (AS-61250-BIOT, Anaspec Inc., Fremont, CA) for 2 hours at 22°C. We washed the beads three times in PBS with 0.1% Tween 20 to remove any unbound antibody. We resuspended the antibody-coated beads from the final wash in BRB80 with 1 nM MTBD to approximately 0.05 % (w/v) bead concentration and incubated for 1 hour at 22°C. We washed the beads three times in BRB80 to remove unbound MTBD. We resuspended the MTBD-coated beads from the final wash in tris assay buffer (50 mM Tris-HCl, pH 8, 50 mM potassium acetate, 2 mM MgCl_2_, 1 mM EGTA, 1 mM DTT, 10 μM Taxol, and 10% glycerol) and stored them at 4°C with 10 mg/mL bovine serum albumin (BSA).

### The optical tweezer and its calibration

We performed force-dependent binding affinity assays using an optical tweezer with a 1064 nm, 10 W Ytterbium fiber laser (YLR-10-1064-LP, IPG Photonics, Oxford, MA) focused on the sample plane by an oil immersion objective (CFI60 Plan Apochromat Lambda 60x N.A. 1.4, Nikon Instruments, Inc., Melville, NY), as previously described (49). Multiple polarizing beam splitters, zero-order half-wave plates, beam traps, and neutral optical density filters (ThorLabs, Inc., Newton, NJ) direct, condition, and regulate the laser power at the sample plane. We monitored the power of the trapping laser at the sample plane by measuring the power of the laser before entering the objective with a photodiode sensor (S121C, Thorlabs, Inc., Newton, NJ), which we correlated to the power at the sample plane using a microscope slide thermal power sensor (S175C, Thorlabs, Inc., Newton, NJ) and power meter interface (PM100USB, Thorlabs, Inc., Newton, NJ).

An integrated micropositioner/nanopositioner (MicroStage and Nano-LP100, Mad City Labs Inc., Madison, WI) driven by USB3 controllers (Micro-Drive2 and Nano-Drive3, respectively, Mad City Labs Inc., Madison, WI) positions and focuses the sample on the image plane of the objective. A Grasshopper3 1.5 MP Mono USB3 Vision CCD camera (imaging chip: ICX825, Sony Semiconductor Solutions Corporation, Atsugi, Japan); camera: GS-U3-15S5M-C, FLIR Systems, Inc., Wilsonville, OR) monitors the sample under test. Brightfield Köhler transmission illumination generated by a LuxeonStar 1030 mW Royal-Blue (448 nm) Rebel LED on a SinkPAD-II 20 mm Star Base (SP-01-V4, Quadica Developments Inc., Lethbridge, Canada) through a series of optical (lenses and irises) and supporting mechanical condenser elements (Thorlabs, Inc., Newton, NJ) and a high NA oil condenser (T-C HNA-OIL High N.A. DIC Lens, Oil N.A. 1.40, Nikon Instruments, Inc., Melville, NY) illuminates the beads, and fluorescence excitation generated by an LED light source (DC2200, Thorlabs, Inc., Newton, NJ) and filtered through a TRITC/Cy3/TagRFP/AlexaFluor 546 filter set (39004, Chroma Technology Corporation, Rockingham, VT) allows us to visualize the fluorescently-labeled polarity-marked microtubules. We detect the displacement of the bead from the trap center using quadrant photodiode-based (QPD, QP45-Q-HVSD, First Sensor, AG, Berlin, Germany) back focal plane detection (50).

We used power spectral density analysis on the position of the trapped bead to calibrate the trap stiffness for each bead (0.09-0.1 pN/nm) (51). Briefly, we converted the QPD voltage into the displacement of the bead from the trap center using the position detection sensitivity factor (β in V/μm) obtained from a stuck bead calibration (51). Then, we divided the measured voltage signal β and multiplied the resultant by trap stiffness to get force traces.

### Force-dependent dissociation assays

We prepared flow cells with channels using piranha-cleaned silanized coverslips (22×22 mm), as previously described (52). We washed the flow channel with BRB80 and then incubated 0.2 mg/mL anti-rhodamine (TRITC) antibody (A-6397, Invitrogen, Waltham, MA) in it for 5 minutes. We washed out excess antibody with BRB80 and then incubated the flow channel with 1% Pluronic F-127 (PK-CA707-59000, PromoCell Inc., Heidelberg, Germany) for 5 min to passivate the surface. We washed out excess Pluronic F-127 with BRB80 and then incubated the flow channel with a polarity-marked microtubule solution diluted in BRB80T for 5 min. We washed out excess microtubules with BRB80T. We 20-fold diluted the MTBD-coated beads in Tris-assay buffer with antifade reagents (125 nM glucose oxidase (G2133, Sigma-Aldrich, St. Louis, MO), 32 nM catalase (C9322, Sigma-Aldrich, St. Louis, MO), 40 mM D-glucose (VWRV0188, VWR Life Science, Solon, OH), 1% β-mercaptoethanol (VWRVM131, VWR Life Science, Solon, OH)) and flowed them into the channel. We sealed the flow channel with nail polish to prevent the evaporation of the solvent.

We trapped an MTBD-conjugated bead and brought it to approximately 50 nm above the plus-end extension of an immobilized polarity-marked microtubule with the optical tweezer. We oscillated the nanopositioning stage in a triangle-wave pattern (2 μm amplitude, 0.2 Hz) with a custom-written LabVIEW virtual instrument (VI, NI, Austin, TX). Given the trap stiffness of 0.09-0.1 pN/nm, this triangle-wave pattern stage movement would correspond to a loading rate of 145-160 pN/s, assuming negligible stretching in the MTBD-antibody-biotin-streptavidin-bead system. However, we found that the actual loading rate was 40 pN/s when considering the system's overall compliance (Supporting Material). We observed binding/unbinding events for approximately 30% of MTBD-coated beads, suggesting that most detected interactions were due to single-molecule binding and unbinding events (53, 54). We tested 10-15 beads in all cases. We found that some beads showed non-specific binding to the coverslip in the absence of microtubule. Therefore, we checked every bead in the presence and absence of microtubules and excluded data collected from beads that showed non-specific binding to the coverslip.

### Analysis of optical tweezer-based force-dependent dissociation experiments

We acquired the optical tweezer data (force traces) at 1 kHz using an FPGA (PXI7854R, NI, Austin, TX) and custom-written LabView VIs. We filtered the data using a Savitzky-Golay filter (polynomial order 4 and frame length 61 ms) (55). When a microtubule-binding event occurred, the moving stage pulled the bead from the trap's center, causing a peak in the force trace.

We distinguished binding/unbinding events from the noise using custom-written MATLAB (The MathWorks, Inc., Natick, MA) scripts. To distinguish binding events from the noise, we calculated the standard deviation of each force trace and took an initial pass at identifying binding/unbinding events by finding the peaks in the force-spectroscopy data that are greater than four standard deviations above the mean force. This first pass overestimates the noise because the calculation includes the signal. Therefore, we excluded the identified peaks from the time-series data and calculated the standard deviation of the reduced data, i.e., the standard deviation of the noise. We took a second pass at identifying binding/unbinding events by finding the peaks that were at least 4.5 standard deviations above or below the mean. We repeated this process iteratively until we found no additional peaks of 4.5 standard deviations above the noise. We set the force values of 4.5 standard deviations above and below the mean as the upper and lower thresholds, respectively, and we take any data that surpass these thresholds as an identified binding event.

We calculated the bound time for each identified event as the interval between the time that the force trace last crossed the 1 standard deviation above or below the mean level (the upper and lower transversal force, respectively) before the peak data point and the time that the force trace next passed the transversal force after the peak data point (Fig. S4 *C* in the Supporting Material for example peaks with upper and lower threshold forces, and upper and lower transversal forces shown). We used the peak force as the unbinding force associated with the event.

We further analyzed each peak to identify spurious events that made it through our peak identification process. We excluded data points that fall into one of two categories, low-force long-duration and high-force double-peak events (Fig. S4 *C*), because neither of these types of events represents single MTBD-microtubule unbinding events (see Supporting Material for details).

We plotted the cumulative distribution function of the bound times, CDF(bound time), for the MTBDs in the force-dependent dissociation assays and fit the CDFs with an exponential function (56)

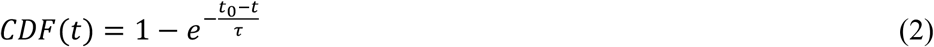

where *t* is the bound time for each unbind event, t_0_ is the minimum resolvable binding time (57), and τ is the characteristic force-dependent unbinding time.

## Results

To investigate the role of long-range electrostatic interactions in regulating the binding affinity of the dynein MTBD for microtubules, we performed binding affinity and dissociation assays using an SRS-domain high-binding affinity locked mouse cytoplasmic dynein MTBD (Materials and Methods) (22). We compared results from the wild-type MTBD to MTBD mutants with charge-altering amino acid substitutions that were remote from the MTBD-microtubule-binding interface under multiple charge-screening conditions (Materials and Methods) (22). We made these changes far from the microtubule-binding interface (D3402A and E3320A, Fig. 1 *B*) to alter long-range electrostatic interactions (34) but not affect short-range interactions (salt bridges and hydrogen bonds) at the binding interface (29), alter dynamic salt bridges (24) or cause structural changes.

Point mutations can alter the structural stability of proteins depending on the location and type of the mutation. Such structural changes could affect the binding affinity of a protein for its partner biomolecule or ligand (58, 59) in ways that are not strongly correlated with long-range electrostatic interactions. Therefore, we assessed the effect of the mutation on the stability of the D3402A and E3320A mutants using circular dichroism (Fig. S2 *A*). We found no significant difference between the ɑ-helical content of the wild-type and the D3402A mutant MTBDs (p-value > 0.05, Fig. S2 *B*, two-sample t-test). However, the ɑ-helical content E3320A was reduced 0.4-fold as compared to WT-MTBD (p-value < 0.001, Fig. S2 *B*, two-sample t-test).

We further assessed the impact of the D3402A and E3320A charge-altering mutations on MTBD protein structure by performing stability calculations using SDM (41) and DynaMut (40). In accordance with the CD results (Fig. S2 *A*), both algorithms predicted that the E3320A mutation should destabilize and the D3402A mutation should stabilize the MTBD’s structure (see Table S1, Fig. S2 *B*, and accompanying text in the Supporting Material for details).

Together, the CD and computational algorithms both predicted that the E3320A mutation caused significant structural differences. It was impossible to deconvolve any changes in binding affinity due to the E3320A mutation from differences due to the long-range electrostatic changes. Therefore, we only used the WT and D3402A mutant proteins in our experiments to probe the effect of long-range electrostatic interactions on dynein MTBD-microtubule binding.

### The binding affinity of D3402A mutant MTBDs for the microtubule is higher than wild-type MTBDs

We measured the dissociation constant, *K*_d_, of the MTBDs and microtubules using microscale thermophoresis (MST, Materials and Methods) to quantify the effect of long-range electrostatic interactions on the binding affinities of MTBD for microtubules. We titrated increasing concentrations of microtubules into a solution containing 50 nM fluorescently-labeled MTBD, acquired MST traces, and found that the normalized fluorescence of MST traces decayed slower and to a higher steady-state normalized fluorescence with increasing concentration of microtubules (Fig. 1 *C*). These data suggest that MTBD bound to microtubules has lower thermophoretic mobility than free MTBD.

We fit the MST traces to a linearized model of MST (Materials and Methods) to calculate the fraction of MTBD bound to microtubules and plotted the results as a function of microtubule concentration (Fig. 1 *D*). Then, we fit Hill–Langmuir binding curves (Materials and Methods) for the WT and the D3402A mutant, and we found that *K*_d_ is 11.5-fold higher (p-value = 0.0014, two-sample t-test) for WT than for D3402A MTBD constructs (*K*_d_ = 4.48 ± 0.29 μM and *K*_d_ = 0.39 ± 0.06 μM, mean ± standard error, N = 3, respectively) in the absence of additional salt (Fig. 1 *D*).

The MST results show that the binding affinity of the D3402A mutant MTBDs for the microtubule is higher than wild-type MTBDs, suggesting that long-range electrostatic interactions either guide the MTBD’s binding to the microtubule, stabilize the MTBD-microtubule bound structure, or both.

### The force-dependent unbinding rate of D3402A mutant MTBDs from microtubules is slower than wild-type MTBDs

External forces can regulate the biophysical properties of dynein by altering the binding affinity of MTBD towards microtubules depending on the direction of applied force (54, 60). We measured the force magnitude and direction-dependent MTBD-microtubule dissociation rate using optical tweezers (Materials and Methods) to study the effect of long-range electrostatic interactions on the force-dependent MTBD-microtubule interactions. We conjugated the MTBD to beads (Materials and Methods) and presented them to polarity-marked microtubules (Materials and Methods) by oscillating the stage position (triangle wave of displacement) with respect to the optical trap (Materials and Methods, Fig. 2 *A*). We captured time-series force spectroscopy data of MTBD-microtubule interactions (Fig. 2 *B* and 2 *C*), binned binding events as occurring under hindering (microtubule plus-end directed) and assisting (microtubule minus-end directed) loads (Fig. 2 *A*), and found the bound time and unbinding force for each binding event (Fig. 2 *C*, Materials and Methods).

**Figure 2.**
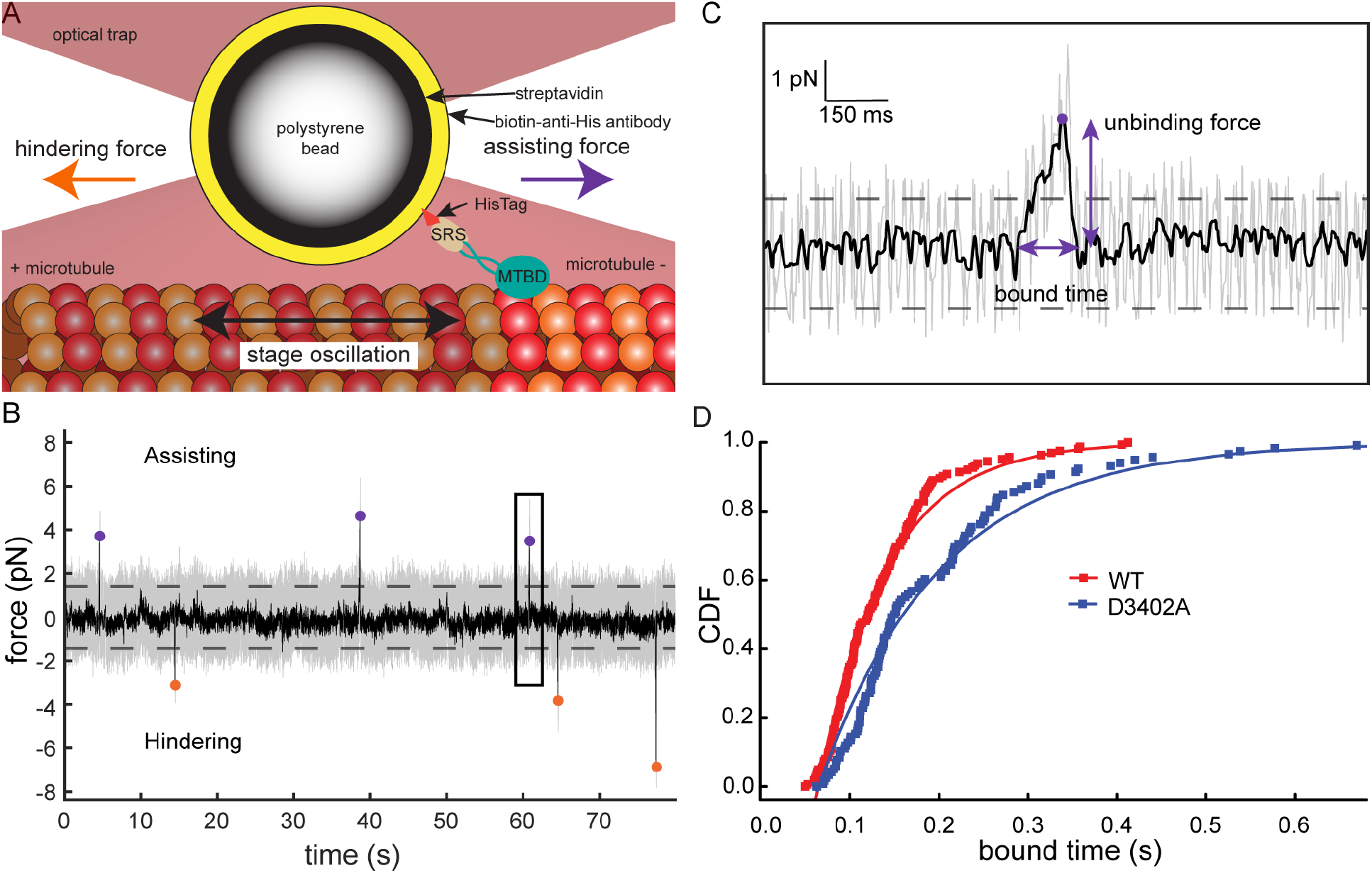
The force-dependent MTBD-microtubule dissociation rate measured using optical tweezers is slower for the D3402A mutant than wild-type MTBDs. (*A*) Schematic representation of force-dependent dissociation assays using optical tweezers. The SRS-MTBD construct binds to a streptavidin-coated polystyrene bead through a biotinylated anti-His Tag antibody that recognizes the C-terminal 6x His domain. The laser (*red*) traps the bead, and the MTBD transiently binds to polarity marked microtubules (minus end GMPCPP stabilized seeds (*bright*) and Taxol-stabilized plus end extensions (*dim*)). The oscillating triangle wave of stage displacement (*black arrow*) causes increasing assisting (*purple arrow*) or hindering (*orange arrow*) forces on the microtubule bound MTBD. (*B*)The MTBD-microtubule-binding detecting algorithm (Materials and Methods) detects events under assisting (*purple dots*) and hindering (*orange dots*) that are greater than 4.5 standard deviations of the noise (*dashed lines*) above the mean in a typical filtered optical tweezers force-dependent assay trace. (*C*) Detail of the boxed binding event in (*B*) indicating the bound time and unbinding force (*purple arrows*). The raw (*gray*) and filtered (*black*) force spectroscopy traces are shown in both (*B*) and (*C*). (*D*) Cumulative distributions of bound times (*squares*) for wild-type (*red*) and D3402A mutant (*blue*) MTBDs on microtubules. We acquired the bound time data by subjecting MTBDs bound to microtubules to increasing force-ramp loads at a constant loading rate of approximately 40 pN/s (Materials and Methods and Supporting Material). The CDFs were statistically different from each other (p-value < 0.001, two-sample K–S test). We fit these data to an exponential cumulative distribution function (Eq. 2) with *t*_0_ = 0.065 s (*lines*) and found that the characteristic unbinding time was τ = 0.076 ± 0.001 s (fit parameter ± SE of the fit, N=163) for wild-type MTBDs and τ = 0.137 ± 0.002 s (N=117) for the D3402A mutant MTBDs.

We plotted the cumulative distribution of single-molecule binding event bound times for the wild-type MTBD and the D3402A mutant MTBD, and we fit the data to exponential cumulative distribution functions (CDF, Eq. 2) for events subject to assisting and hindering loads (Fig. S5 in the Supporting Material). We found that, while the binding time was a function of the mutation (Fig. S5), the CDFs were not statistically different from each other (p-value > 0.05 in all cases, two-sample Kolmogorov–Smirnov test) as a function of load direction. These results indicate that the force-dependent unbinding of the MTBD from the microtubule is symmetric with respect to the direction of load, which is consistent with previous reports (54). Therefore, we pooled the assisting and hindering data for further analysis.

We compared the pooled distributions of bound time for the wild-type and D3402A mutant MTBD-microtubule force-dependent dissociation events, and we found that they came from statistically distinct populations (Fig. 2 *D*, p-value < 0.001, two-sample K-S test). We fit the exponential cumulative distribution function (CDF, Eq. 2) to the cumulative distribution of bound times for the wild-type MTBD and D3402A mutant MTBD-microtubule force-dependent dissociation events (Fig. 2 *D*). We found that the characteristic bound time, τ, was approximately two-fold larger for the D3402A mutant than wild-type MTBDs (Fig. 2 *D*).

The force-dependent results show that the unbinding rate of D3402A mutant MTBDs from microtubules is slower than wild-type MTBDs, suggesting that long-range electrostatic interactions propagate through the MTBD and stabilize short-range salt bridges at the MTBD-microtubule binding interface.

### Increasing the ionic strength of the solution differentially affects the MST-derived dissociation constant and the optical tweezer determined force-dependent unbinding rate

We repeated the MST and optical tweezer assays on the wild-type and the D3402A mutant MTBDs in the presence of additional salt to decrease the Debye length in the solvent from approximately 0.8 nm for no additional salt to 0.6 and 0.4 nm for an additional 100 mM and 300 mM KCl, respectively (Eq. 1). Additional salt increases the charge screening of long-range electrostatic interactions that act through the solvent.

We found that the addition of 100 mM KCl had a small effect (only a 31% reduction, p-value = 0.03, two-sample t-test, Fig. 3 *A*) on the *K*_d_ for wild-type MTBDs, as measured using MST (Materials and Methods). However, the addition of 100 mM KCl had a significantly greater effect (a 14-fold increase, p-value < 0.001, two-sample t-test, Fig. 3 *A*) on the *K*_d_ for the D3402A mutant MTBDs than wild-type MTBDs. The additional salt reduced the 11.5-fold difference in *K*_d_ between the wild-type MTBDs and D3402A mutant MTBDs in the absence of additional salt to only a 1.8-fold difference (p-value < 0.001, two-sample t-test, Fig. 3 *A*), and *K*_d_ for wild-type MTBDs in the absence of additional salt was nearly the same as for the D3402A mutant with 100 mM additional KCl (4.48 ± 0.29 μM, and 5.49 ± 0.12 μM, p-value = 0.14, two-sample t-test, Fig. 3 *A*).

**Figure 3:**
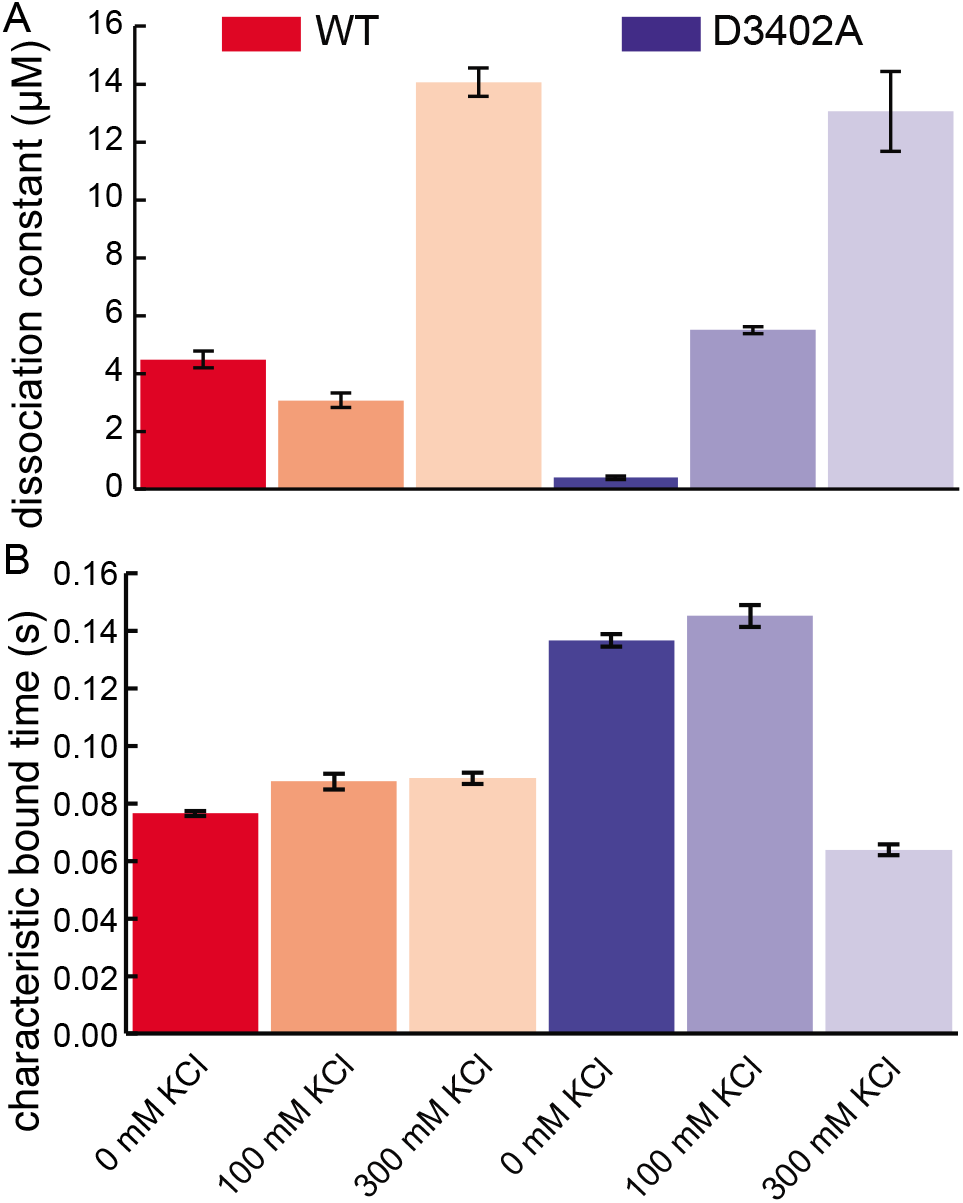
Solution ionic strength differentially affects the dissociation constant and characteristic bound times of wild-type and D3402A mutant MTBDs on microtubules. (*A*) The dissociation constant, *K*_d_, for the wild-type (WT) and D3402A mutant MTBDs obtained by fitting Hill–Langmuir binding curves to the fraction bound for each KCl concentration case in MST experiments. Error bars represent the standard error of the mean (N=3 in each case) from the MST-derived fits. The fluorescently-labeled MTBD concentration was 50 nM, and the concentration of microtubules varied from 0 to 23 μM in all MST experiments, similar to Figure 2 *B*. (*B*) Characteristic force-dependent unbinding time, *τ*, obtained from the fits of cumulative distributions of bound times data to Eq. 2 with *t*_0_ = 0.065 s. Error bars represent standard errors of the fits in both panels. MTBD construct and additional salt concentrations legends and labels apply to both panels.

Adding 200 mM more salt in MST, to a concentration of 300 mM additional KCl, significantly increased the *K*_d_ for both wild-type MTBD and the D3402A mutant MTBD, from 3.07 ± 0.25 μM to 14.6 ± 0.49 μM (p-value < 0.001, two-sample t-test, Fig. 3 *A*) and from 5.49 ± 0.12 μM to 13.0 ± 1.4 μM (p-value = 0.02, two-sample t-test, Fig. 3 *A*), respectively. The additional salt at 300 mM KCl further masked the differences between the *K*_d_ for wild-type MTBDs and the D3402A mutant MTBDs (14.6 ± 0.49 μM and 13.0 ± 1.4 μM, p-value = 0.40, two-sample t-test, Fig. 3 *A*).

We also found that the addition of salt (both 100 mM and 300 mM KCl) had a statistically negligible effect (p-values > 0.05, two-sample K-S tests, Table S3) on the cumulative distribution of bound times for wild-type MTBDs when subject to increasing load (approximately 40 pN/s, Supporting Material) by examining the optical tweezer. Additionally, the addition of 100 mM KCl had a statistically negligible effect (p-value = 0.77, two-sample K-S test, Table S3) on the cumulative distribution of bound times for D3402A mutant MTBDs when subject to increasing. Furthermore, the addition of 300 mM KCl restored the cumulative distribution of bound times for D3402A mutant MTBDs to being statistically indistinguishable from the wild-type MTBDs (p-value > 0.05, two-sample K-S tests, Table S3).

We also fit the exponential cumulative distribution function (CDF, Eq. 2) to the cumulative distribution of bound times for the wild-type MTBD and D3402A mutant MTBD-microtubule force-dependent dissociation events in 100 and 300 mM KCl additional salt. We found that the characteristic bound time, τ, was nearly 1.75-fold larger (approximately 0.14 s, based on the mean of τ for 0 and 100 mM additional KCl, Fig. 3 *B*) for the D3402A mutant MTBDs at no and low (100 mM) additional salt, as compared to wild-type MTBDs (approximately 0.08 s, based on the mean of τ for all three conditions, Fig. 3 *B*). Additionally, we found that the highest (300 mM) additional salt concentration essentially masked the differences between the characteristic bound time of D3402A and wild-type MTBDs (0.8-fold, but p-values > 0.05 from all two-sample K-S tests, Table S3).

Together, these results show that dissociation constant and force-dependent unbinding rate differentially depend on the ionic strength of the solution. These results suggest that long-range electrostatic interactions are more strongly attenuated by salt at thermodynamic binding equilibrium than when bound to the microtubule.

## Discussion

In the present study, we investigated whether long-range electrostatic interactions, i.e., electrostatic interactions between charged amino acids not at the immediate binding interface, play a significant role in binding cytoplasmic dynein microtubule-binding domains locked into the high-binding affinity state to microtubules. We used both site-directed mutagenesis (to change a charge remote from the binding interface, Fig. 1 *B*, but did not change the structure, Fig. S2) and salt-induced charge screening (to change the Debye length, Eq. 1) to probe these effects. We found that long-range electrostatic interactions do play a significant role in the binding affinity by affecting both the dissociation constant, *K*_d_, and the unbinding rate, *k*_off_.

Using microscale thermophoresis, we found that the dissociation constant (*K*_d_) of the wild-type MTBD was more than 10-fold higher than the positive charge neutralizing D3402A mutant (Fig. 1 *D*). This result strongly suggests that long-range electrostatic interactions play a critical role in the binding affinity, as the D3402A mutation is 2.4 nm from the binding interface.

We further probed the effect of long-range electrostatic interactions by increasing the electrolyte concentration in the solution. We found that the addition of 100 mM KCl did not strongly affect the K_d_ of the wild-type MTBD (Fig. 3 *A*), but it essentially restored the wild-type affinity to the D3402A mutant MTBD (Fig. 3 *A*). The further addition of salt to 300 mM of additional KCl masked the affinity difference of both MTBDs towards microtubules.

Together, the MST results must be understood in the context of the differences between the effects of charge screening on the electric field lines that pass through the solution and those that pass through the solvent-excluded volume of the protein. Debye-Hückel theory (61) shows that the electric potential due to charged amino acid residues at a protein surface decreases by a factor of 1/e (approximately 63% attenuated) each Debye length into the solvent (Eq. 1). In both cells and biological buffers with an electrolyte concentration of approximately 150 mM, the Debye length is approximately 0.8 nm (Eq. 1). Thus, the strength of the electrostatic field lines between aspartic acid 3402 in the MTBD and the microtubule surface that pass through the solvent are attenuated by about 95%. However, many of the electrostatic field lines between D3402 and the microtubule surface pass through the solvent-excluded protein-filled regions of space occupied by the rest of the MTBD. Debye-Hückel theory does not apply in these regions. Thus the magnitude of the electric field strength is unattenuated by charge screening. Moreover, the electric field strength, 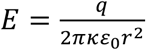 where *q* is the charge on the amino acid, *r* is the distance between the charged amino acid and the microtubule, and *ε*_0_ is the electrical permittivity of free space, is stronger in protein-filled regions of space than solvent-filled regions of space because the dielectric constant, *κ*, is lower in the protein (often assumed to be 4) than in the solvent, which has *κ* ≈ 80 (62, 63). Thus, despite the relatively high concentration of electrolytes, long-range electrostatic interactions can significantly affect binding affinity.

Using optical tweezers to characterize the force-dependent dissociation of the MTBDs from microtubules, we found that the characteristic force-dependent unbinding time when subject to a constantly increasing load, τ, of the positive charge neutralizing D3402A mutant was approximately 1.75-fold higher than the wild-type MTBD (Fig. 2 *D*). We further probed the effect of long-range electrostatic interactions by increasing the electrolyte concentration in solution, and we found that neither the addition of 100 mM nor 300 mM KCl strongly affected τ of the wild-type MTBD (Fig. 3 *B*). Only the addition of 300 mM KCl, but not 100 mM KCl, restored the wild-type value of τ to the D3402A mutant MTBD (Fig. 3 *B*). These results suggest that increasing the long-range electrostatic forces significantly stabilized the MTBD-microtubule bound structure. Moreover, because it took a large amount of added salt to screen the increased long-range electrostatic interactions due to the D3402A mutation, these results suggest that the most significant contribution of the long-range electrostatic interactions in the MTBD-microtubule interface act through regions of space that are minimally solvent accessible, i.e., through the globular MTBD domain. These results are consistent with other systems in which long-range interactions are essential for binding, e.g., TEM1 β-lactamase and its protein inhibitor BLIP (64) and kinesin and microtubules (65).

Taken together, the MST and force-dependent dissociation results show that the charge screening affects the equilibrium dissociation constant (*K*_d_) more strongly, and with more sensitivity (lower salt is necessary) than the unbinding kinetics (*k*_off_, which scales like 1/τ,) of MTBD-microtubule interactions. This suggests that long-range electrostatic interactions are particularly important to aspects of biomolecular bonding systems that are not highly susceptible to charge screening by soluble ions. For example, the solvent and soluble ions are excluded from the MTBD-microtubule interface when the MTBD is bound to the microtubule, which reduces the Debye effect (Fig. 4 *A*, left) and thus the relative insensitivity of the unbinding kinetics (*k*_off_) to additional salt. However, the solvent and soluble ions fill the space between the MTBD and the microtubule when the MTBD is not bound to the microtubule, which screens charges due to the Debye effect (Fig. 4 *A*, right) and thus the relative sensitivity of the equilibrium dissociation constant (*K*_d_) to additional salt.

**Figure 4.**
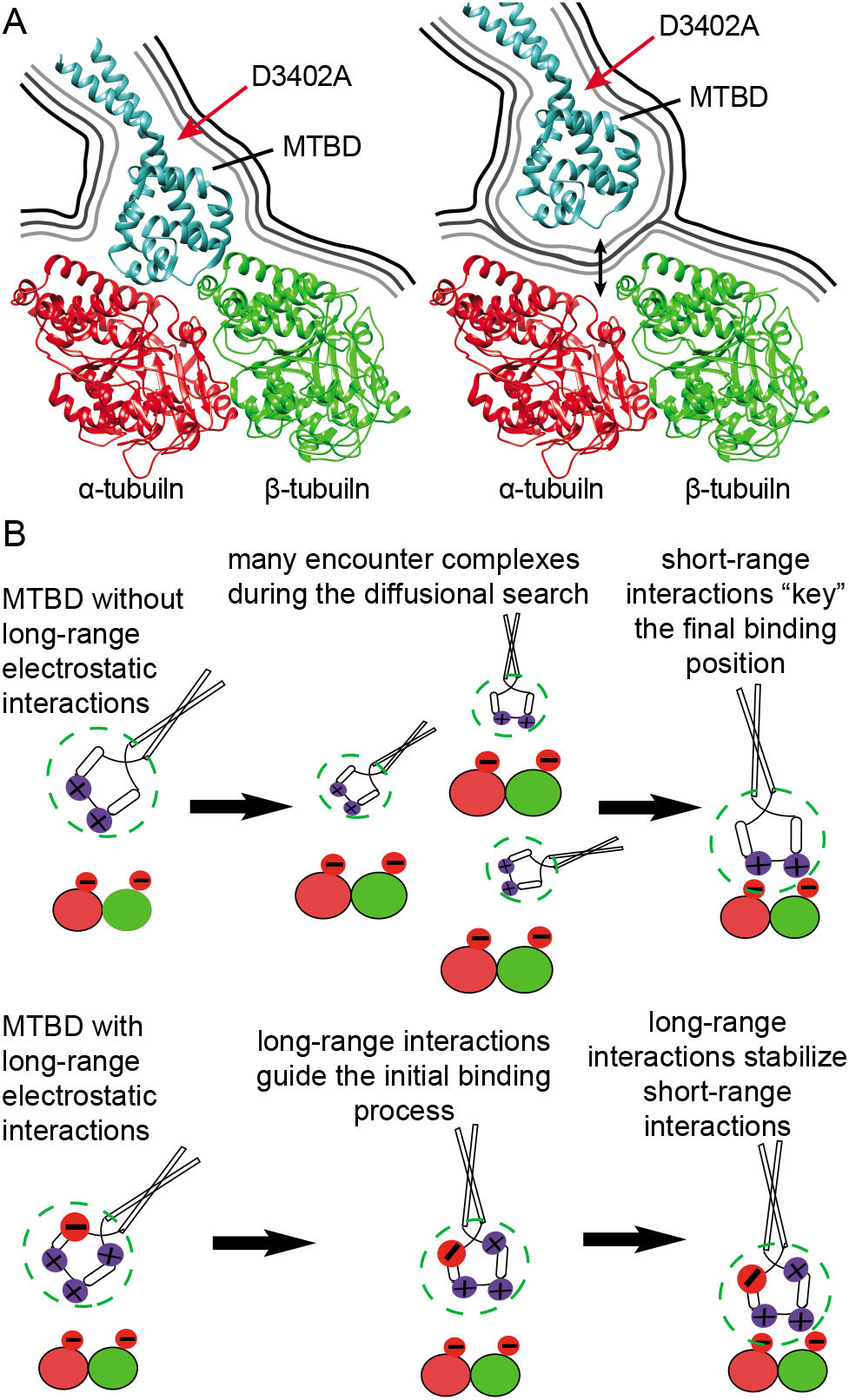
Long-range electrostatic interactions and the Debye effect contribute significantly to the binding of dynein to microtubules. (*A*) Ribbon diagram representation of the MTBD (*teal*) bound to (*left*) and unbound from (*black double-headed arrow*, *right*) a tubulin dimer (PDB ID: 3J1T (24)), similar to Fig. 1 *B*. Approximate Debye length, as it propagates into the solvent-accessible space around the proteins, is shown for 0 mM additional KCl (*dark gray line*), 100 mM additional KCl (*medium gray line*), and 100 mM additional KCl (*light gray line*). (*B*) Model of how long-range and short-range electrostatic interactions mediate MTBD binding to microtubules. In the absence of long-range electrostatic interactions (*top*), there are multiple possible MTBD-microtubule encounter complexes. Long-range electrostatic interactions (*bottom*) provide orientation to initial binding of MTBD to microtubules through electrostatic steering effects and additional stabilization of the bound complex.

Moreover, this model has implications for the molecular details of dynein motility and processivity. It suggests that as an MTBD moves through the electrostatic binding funnel, which guides and aligns MTBDs to the binding interface on the microtubule (34, 35, 65, 66) as well as other biomacromolecular interactions (67–71), it displaces more solvent and reduces the Debye effect, effectively reducing the attenuation of the electrostatic field strength and increasing the electrostatic forces. Thus, this model suggests that a positive physical force feedback mechanism for biomolecular binding driven by long-range electrostatic interactions is a significant contributor to the processivity of dynein motor proteins.

The observed significant decrease in the 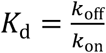 upon changing the charge of an amino acid remote from the binding interface suggests that either the binding kinetics, *k*_on_, increased or the unbinding kinetics, *k*_off_, decreased, or both. The observed increase in τ suggests a decrease in *k*_off_. However, the magnitude of the increase in τ (1.75-fold) is unlikely to explain the magnitude of the decrease in *K*_d_ (11.5-fold). This further suggests that the change in the long-range electrostatic interactions induced by the charge-altering mutation remote from the binding interface increased *k*_on_, and thus the MTBD’s microtubule-binding funnel.

Altogether, our results support our previous computational studies (34, 35), which also highlighted the significant role of long-range electrostatic interactions in guiding the MTBD binding to microtubules by providing an electrostatic binding funnel. Similar long-range electrostatic interaction-guided regulation mechanisms were reported earlier in the case of kinesin motors (65, 66, 72). Together with the results from previous experimental work detailing how the short-range electrostatic interactions that constitute surface charge complementarity at the MTBD-microtubule binding interface (22, 29, 31) and specific, dynamic salt-bridges within the MTBD (24) underly the microtubule-binding affinity, our results help build a more complete picture of how electrostatics govern the molecular mechanics of dynein motility. We propose that long-range electrostatic interactions, short-range electrostatic interactions, and dynamic salt bridges significantly contribute to various aspects of the binding affinity mechanism as a function of the distance the MTBD is from the microtubule. The net charge of the MTBD, which includes charges at and far from the binding pocket, guides the initial binding process (Fig. 4 *B*). The distribution of charge on the MTBD, including long-range interactions remote from the binding interface, can apply electrostatic torques to the MTBD that may be particularly important to reorienting (34) the MTBD as it approaches the microtubules. As dynein’s MTBD achieves a favorable binding pose, our model suggests that the binding kinetics accelerate due to solvent exclusion induced electrostatic charge screening reduction, and once bound, the short-range electrostatic interactions that “key” the alignment (34) and dynamic salt-bridges that regulate affinity can be further stabilized or destabilized by long-range interactions within the MTBD (Fig. 4 *B*). All the effects described in this model are further regulated by the role that the negatively charged, intrinsically disordered, highly posttranslationally modified, C-terminal tail of tubulin plays in guiding the binding process and regulating the bound complex (35).

Beyond the role of electrostatic interactions, acidic amino acid residues, in particular aspartic acid, have been shown to be the primary feature of proteins that stabilizes and solubilizes them in the presence of the high salt concentrations experienced by halophiles, which are extremophiles living in environments with more than five times the salinity of sea water (73). Aspartic acids provide favorable sites for water molecules in the ordered hydration layer to hydrogen bond with the protein surface at higher salt concentration, leading to increased hydration shell stability of aspartic acid-rich protein surfaces (74, 75). By mutating aspartic acid 3402 to an alanine, we could have made the MTBD more susceptible to increased salt concentration, though the additional 300 mM (at the most) KCl is significantly lower than the 2.5-5 M salt concentrations experienced by halophiles. This effect could help explain the difference in the change in the *K*_d_ values in the D3402A mutant MTBD as compared to that in the wild-type MTBD when going from no salt to 100 mM salt (Fig. 3 *A*). Along the same lines, the microtubule-binding dynamics of the wild-type MTBD were significantly less susceptible to the increased salt concentration, as all three conditions considered showed similar unbinding rates (Fig. 3 *B*). This analysis reiterates that the physical effects of the amino acids in proteins act at multiple levels, with multiple modes, and across multiple scales.

These results, and the models they suggest, have significant implications on our understanding of processivity in the dynein family of motor proteins. All dynein family members likely rely on specific short-range electrostatic interactions and the ability to change these interactions as a function of mechanochemical cycle state, coiled-coil stalk registration, and MTBD structural conformation. However, our results suggest non-specific long-range electrostatic interactions that provide bound-state stabilization may be more prominent in processive cytoplasmic dyneins than axonemal dyneins. Moreover, tuning the long-range electrostatic interactions without altering the binding interface residues could be a useful strategy to engineer the motor proteins with different processivities. Finally, our results also suggest that similar regulation mechanisms may be important in other motor proteins and for other protein-protein interactions in general.

## Author Contributions

J.A. designed research; A.Pab., L.C., M.S., and J.E. performed research; A.G., A.Pau., and J.A. contributed analytic tools; A.Pab., L.C., S.G., A.Pau., A.G., and J.A. analyzed data; A.Pab. and J.A. wrote the manuscript.

## Acknowledgments

We thank Lin Li and Emil Alexov for insightful discussions and collaboration related to the hypothesis of the present work. We also thank Jonathon Howard and the Howard Lab for the tubulin, Ron Vale for the MTBD plasmid (Addgene plasmid # 22380), Hugo Sanabria for the use of his equipment, Robert Latour and the Clemson Biomaterials Engineering and Testing Core Instruments for the use of the circular dichroism spectropolarimeter, and the entire Alper Lab for fruitful discussions. We also thank Marija Zanic for helpful feedback and fruitful discussions.

This work was supported in part by Clemson University’s R-Initiative CU-MRI grant for the microscale thermophoresis instrument, the Department of Physics and Astronomy, the College of Science, the Creative Inquiry Program, and the Clemson University Professional Internship and Co-op Program (UPIC). Additional support was provided by the National Institute of Allergy and Infectious Diseases (NIAID) of the National Institutes of Health under award number R15AI137979 and the National Institute of General Medical Sciences (NIGMS) of the National Institutes of Health under award number P20GM109094.

The authors declare that there are no conflicts of interest.

## Supporting References

References (76–78) appear in the Supporting Material.

