## Supporting Material for "Long-Range Electrostatic Interactions Significantly Modulate the Affinity of Dynein for Microtubules"

#### Supporting Methods

*Quantification of MTBD expression purity*: We used densitometry of Coomassie blue stained SDS-PAGE gel electrophoresis. We stained the polyacrylamide protein gels (Fig. S1) with a Coomassie G-250 dye-based reagent and a rapid staining procedure, per the manufacturer's instructions (24594, GelCode Blue Safe Protein Stain, Thermo Scientific, Waltham, MA). We scanned the gels (B11B178011, Epson Perfection V700 Photo, Epson, Inc., Hillsboro, OR) and performed quantitative densitometry using ImageJ (1, 2). To quantify the purity of the purified MTBD proteins (Fig. S1, *top*), we made a serial dilution of the HisTrap HP column-purified, PD-10 desalted fractions of interest (Fig. S1, *bottom*). We quantified the optical density of each band and identified the highest concentration of MTBD for which the measured density was linear with the amount of protein loaded in the well. We then calculated the fraction of optical density in the entire lane that corresponded to the principal band (62.5 kDa). We found that the WT was 87% pure, D3402A was 91% pure, and E3320A was 93% pure.

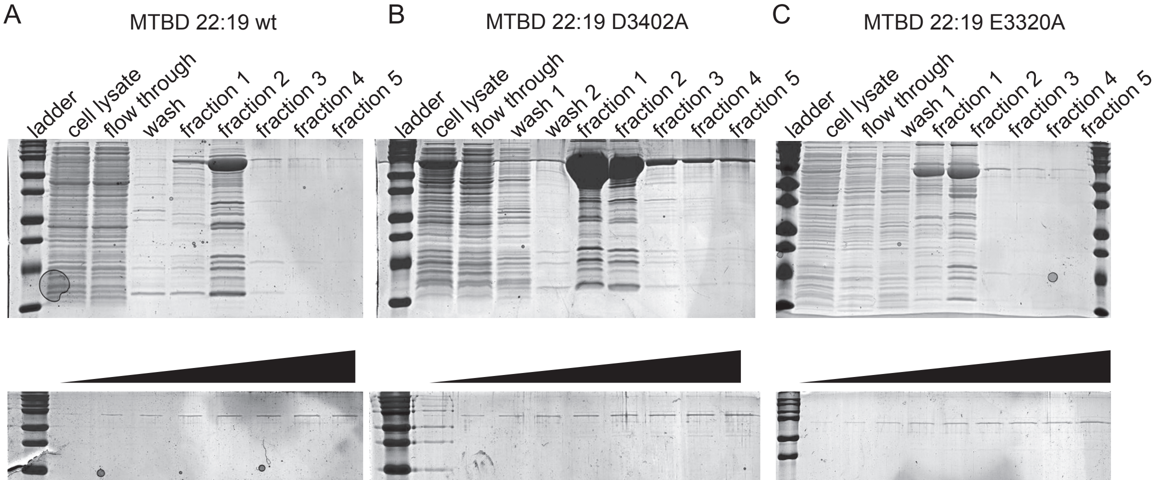


**Figure S1 Quantification of MTBD expression purity.** Gels of aliquots saved from various steps (clarified cell lysate, HisTrap column flow through, HisTrap column wash, and 1 column volume election fractions) from the purification of the (*A*) wildtype, (*B*) D3402A mutant, and the (*C*) D3320A mutant constructs (*top*). Fractions 1 and 2 from each purification were pooled, PD-10 column desalted, and loaded into another gel (*bottom*) at increasing (25 – 250 ng of total protein per well) quantities (*black triangle* to indicate increasing load). Quantitative densitometry indicated 87% purity for wildtype in (*A*), 91% purity for D3402A in (*B*), and 93% purity for E3320A in (*C*).

*Assessment of mutant MTBD structural stability using circular dichroism*: We used circular dichroism (CD) to assess the structural stability of the purified MTBD protein constructs. We recorded CD spectra (206-250 nm) of the purified MTBD proteins with a spectropolarimeter (J-810, Jasco, Inc., Easton, MD) at 25 °C in PBS buffer containing 10% glycerol using a 1 mm pathlength cuvette (1-Q-1, Starna Cells, Inc., Atascadero, CA). We recorded the CD spectral signal of MTBD constructs using approximately 0.5 mg/mL concentration and reported the molar ellipticity to ensure the signal is corrected for any concentration difference among different samples.

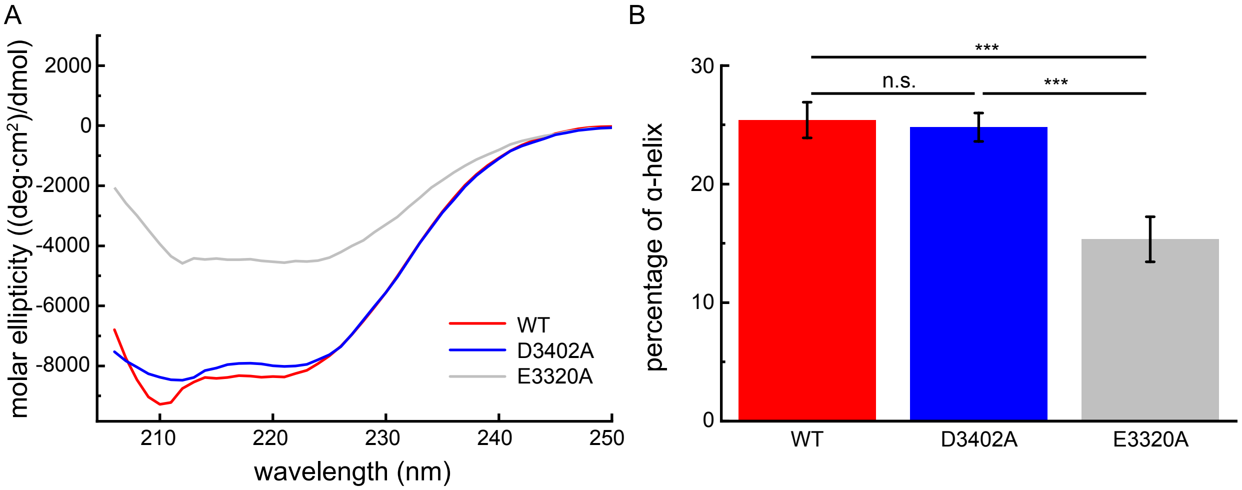


**Figure S2 The D3402A mutation retains the wt-MTBD structure.** (*A*) CD spectra of the wild-type (WT) and mutant (as indicated) MTBD protein constructs. The CD spectra were recorded in PBS buffer at a concentration of 0.5 mg/mL. (*B*) The percentage of α-helical content of wild-type and mutant MTBD constructs. Error bars represent SE and N=2. We found that the difference between α-helical content of WT and D3402A was not significant (p-value > 0.05, n.s., two-sample t-test), but that the difference between α-helical content of E3320A was (p-value < 0.001, ***, two-sample t-test) in each case.

We assessed the effect of mutation using the percentage of calculated α-helical content of the constructs as a proxy for the stability of the designed MTBD mutants (D3402A and E3320A). We estimated the fraction of helical content (*FH*) of the MTBDs, where

$$FH = 0.00514A + 0.00297 \qquad \text{(S1)}$$

and A is the slope of the CD spectrum (molar ellipticity/nm) between 230-240 nm in (deg·cm$^2$)/(dmol·nm) (3). We found no significant difference between the α-helical content of wildtype and the D3402A mutant (p-value > 0.05, two-tailed t-test, Fig. S2 *B*). However, the α-helical content E3320A was reduced approximately 0.4-fold as compared to the wildtype MTBD (p-value < 0.0001, Fig. S2 *B*).

*Assessment of mutant MTBD structural stability using computational techniques*: We used DynaMut (4) and the Site Directed Mutator (SDM) (5) to predict the change in the folding free energy and thus the stability of the MTBD upon mutation. DynaMut implements normal modal analysis to estimate the effect of mutation on protein stability. SDM calculates the stability score using a statistical potential energy function based on the frequency and identity of amino acid replacements occurring within specific protein structural environments that tend to be tolerated by or destabilize the structure. Both methods calculate the change in the change of the free energy upon folding (ΔΔG) of a protein due to mutation. A negative ΔΔG indicates that the mutation destabilizes and a positive ΔΔG indicates that the mutation stabilizes the structure (4).

We used the high affinity dynein microtubule binding domain - tubulin complex cryo-EM (PDB ID: 3J1T (6)) and details about mutation, i.e., the aspartic acid to alanine mutation for D3402A and the glutamic acid to alanine mutation for E3320A, to predict the stability changes upon mutation. In accordance with the CD results (Fig. S2), both algorithms predicted that the E3320A mutation should destabilize and the D3402A mutation should stabilize the MTBD's structure (Table S1).

**Table S1 Changes in the folding free energy, ΔΔG, associated with the charge-altering mutations.** Summary of changes in the MTBD's folding free energy and stability predictions from DynaMut and SDM.

| MTBD | ΔΔG (kcal/mol) DynaMut | ΔΔG (kcal/mol) SDM |
|---|---|---|
| D3402A | 0.15 (stabilizing) | 1.24 (stabilizing) |
| E3320A | -0.76 (destabilizing) | -0.10 (destabilizing) |

We further investigated the specific interatomic interactions that DynaMut predicts to be changed upon mutation. We found that in both cases multiple bonds are missing from the mutant structures that occur in the wildtype (Fig. S3, *colored arrows*, *left*). However, these may be partially stabilized by the addition of a halogen bond in the case of D3402A,

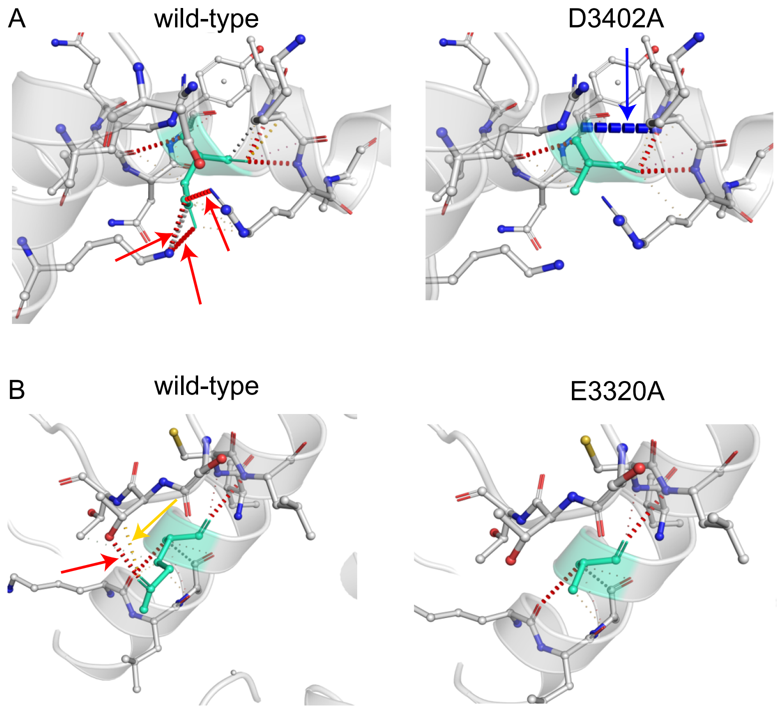


**Figure S3 Point mutations alter the interatomic interactions present in MTBD.** (*A*) A comparison of the interatomic interactions in wild-type (*left*) and the D3402A (*right*) mutant MTBDs obtained from Dynamut. (*B*) A comparison of the interatomic interactions in wild-type (*left*) and E3320A (*right*) mutant

MTBDs obtained from Dynamut. In both panels, the mutated residues are shown in light-green sticks/ribbon diagram highlight. New and broken hydrogen bonds (*red arrows* indicating *red broken lines*), halogen bonds (*blue arrows* indicating *blue broken lines*), and ionic interactions (*yellow arrows* indicating *yellow broken lines*) are shown.

Together, the results showed that the D3402A mutation left the structure essentially unaltered, but that the E3320A mutation caused significant structural adorations. Therefore, we only characterized the binding properties of the D3402A.

*Identification of spurious binding/unbinding events*: Within the harmonic approximation, the restoring force ($F_{\mathrm{trap}}$) experienced by the bead in an optical trap is

$$F_{\mathrm{trap}} = -k_{\mathrm{trap}} x_{\mathrm{bead}} \qquad \text{(S2).}$$

where, $k_{\mathrm{trap}}$ is the stiffness of the trap, and $x_{\mathrm{bead}}$ is the displacement of the bead from the center of the trap. Assuming that the system is a quasi-equilibrium, the force of the trap and the force of the MTBD, $F_{\mathrm{MTBD}}$, are the only two forces exerted on the bead, so

$$F_{\mathrm{trap}} + F_{\mathrm{MTBD}} = 0 \qquad \text{(S3).}$$

Thus

$$F_{\mathrm{MTBD}} = k_{\mathrm{trap}} x_{\mathrm{bead}} \qquad \text{(S4).}$$

is exerted on the MTBD and causes it to undergo force-dependent dissociation (Bell's model). We subjected this system to a triangle wave of stage displacement, and if we make the further assumption that the molecule under test (the SRS-MBTD) and all the linking elements (streptavidin, biotin, anti-His6) are rigid ($k_{\mathrm{MTBD+linkers}} \to \infty$) compared to the trap and that the bead is displaced from the trap immediately upon binding, then we could assume $x_{\mathrm{bead}} = v_{\mathrm{stage}} t_{\mathrm{bound}}$ where $v_{\mathrm{stage}}$ is the stage velocity and $t_{\mathrm{bound}}$ is the bound time

$$F_{\mathrm{MTBD}} = k_{\mathrm{trap}} \left( v_{\mathrm{stage}} t_{\mathrm{bound}} \right) \qquad \text{(S5).}$$

Therefore, the unbinding force ($F_{\mathrm{unbind}}$), which equals $F_{\mathrm{MTBD}}$ at the moment of unbinding, would be a linear function of bound time ($t_{\mathrm{bound}}$) with a slope of $\left( k_{\mathrm{trap}} v_{\mathrm{stage}} \right)$, if all the assumptions are good.

We plotted unbinding force as a function of bound time for each force spectroscopy trace. (e.g., Fig. S4 *A*). We performed a linear regression and identified significant outliers (those points whose standardized residual, $r_i = \frac{e_i}{\mathrm{RSE}\sqrt{1-h_{ii}}}$ where $e_i$ is the residual of the i$^{\mathrm{th}}$ data point, RSE is the residual standard error, and $h_{ii}$ is the leverage of the i$^{\mathrm{th}}$ data point, exceeded a value of 2) from the best fit line (e.g., Fig. S4 *A*, *red circles*) and investigated each of them more closely (e.g., Fig. S4C). We found two types of events that lead to outlying points. Low force, long duration events (e.g., Fig. S4 *C*, *left*) occur when the noise exceeds the 4.5 σ threshold (1 in approximately 300,000 data points) to trigger event detection, but the neighboring measurements do not fall below the upper traversal threshold designating the beginning and end of an event for a long duration (quite rare). These low force, long duration events are likely to be noise mischaracterized as events. High force, double peak events (e.g., Fig. S4 *C*, *right*) occur when a small peak immediately follows a very large one. High force, double peak events are likely to be due to double, rather than, single-molecule binding events because the molecules share the load to yield such a high peak, and the second one fails rapidly once the bead reaches its new equilibrium position in the trap due the remaining bond. We discarded both low force, long duration and high force, double peak events from further analysis.

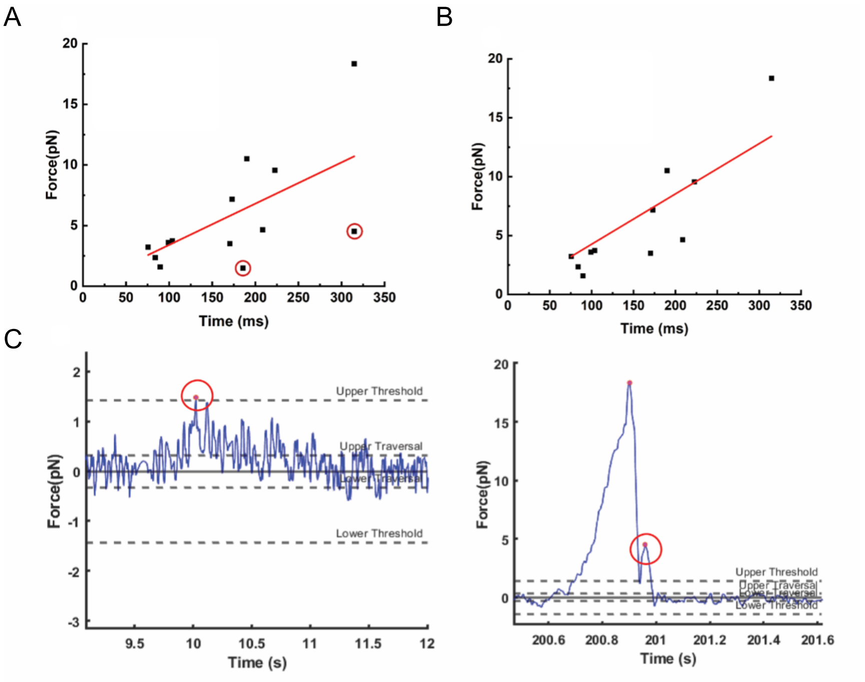


**Figure S4 Unbinding force as a function of bound time plots allow us to exclude the false events.** (*A*) Example unbinding force data plotted as a function of time to unbind (*black squares*) for WT MTBD loaded in assisting direction at a rate of $k_{\mathrm{trap}} v_{\mathrm{stage}} = 144$ pN/s along with the least squares regression line of best fit (*red line*), which has a slope of 34.0 ± 5.3 (fit parameter ±SE of the fit) pN/s, which was significantly (more than four-fold) less than the applied loading rate ($p$ value < 0.0001, two-tailed t-test). Two of the three outliers (*red circles*) are examined in detail in panel C. (*B*) The unbinding force data from (*A*) plotted as a function of bound time (*black squares*) after removing the excluded data along with a least squares regression line of best fit (*red line*). The plot after removing the highlighted data points. The slope obtained from the fitting is 42.6 pN/s. (*C*) Time series force spectroscopy data detailing the outlying data points highlighted in (*A*). Low force, long duration (*left*) and high force, double peak (*right*) events originating as a noise following an unbinding event were ultimately removed from the data set.

*MTBD and linker stiffness*: We re-examined the unbinding force as a function of bound time for the example WT MTBDs data after excluding low force, long duration and high force, double peak events (Figure S4 *C*). We found that the slope was 42.6 ± 4.6 (fit parameter ±SE of the fit) pN/s, which was still significantly (more than three-fold) less than the applied loading rate ($p$ value < 0.0001, two-tailed t-test). This discrepancy is likely due to one or more of the assumptions described above being violated. We suggest that the MTBD and linking elements (streptavidin, biotin, anti-His6, SRS domain) may not be rigid ($k_{\mathrm{MTBD+linkers}} \to \infty$) compared to the trap. If we relax this assumption, then we can model the system as having two finite stiffness springs ($k_{\mathrm{MTBD+linkers}}$ and $k_{\mathrm{trap}}$) in series such that Eq. S5 becomes

$$F_{\mathrm{MTBD}} = k_{\mathrm{eq}}(v_{\mathrm{stage}} t_{\mathrm{bound}}) \quad \text{(S6a)}$$

where $k_{\mathrm{eq}} = \frac{k_{\mathrm{trap}} k_{\mathrm{MTBD+linkers}}}{k_{\mathrm{trap}} + k_{\mathrm{MTBD+linkers}}}$ and thus

$$F_{\mathrm{MTBD}} = v_{\mathrm{stage}} \frac{k_{\mathrm{trap}} k_{\mathrm{MTBD+linkers}}}{k_{\mathrm{trap}} + k_{\mathrm{MTBD+linkers}}} t_{\mathrm{bound}} \quad \text{(S6b).}$$

So, using Eq. S6b and the slope of 42.6 +/- 4.6 pN/s, we estimate that $k_{\mathrm{MTBD+linkers}}$ is 0.038 pN/nm (approximately 40-50% as stiff as the trap), in this case, but it is typical of all the traces.

We further note that for a given equivalent loading rate (~40 pN/s) and 4.5 σ ~ 1.5 pN threshold for event detection yields a minimum resolvable bound time of $t_0$ ~ 40 ms (7).

### Supporting results

*The force dependent MTBD-microtubule dissociation was not a function of force pulling direction*: We fit the data to exponential cumulative distribution functions (CDF, Eq. 2 from the main text) for events subject to assisting and hindering loads (Fig. 2 *A* from the main text) and found that, while the unbinding time was a function of the mutation and the salt concentration, the CDFs were not statistically different from each other (p value > 0.05 in all cases, two-tailed Kolmogorov–Smirnov test).

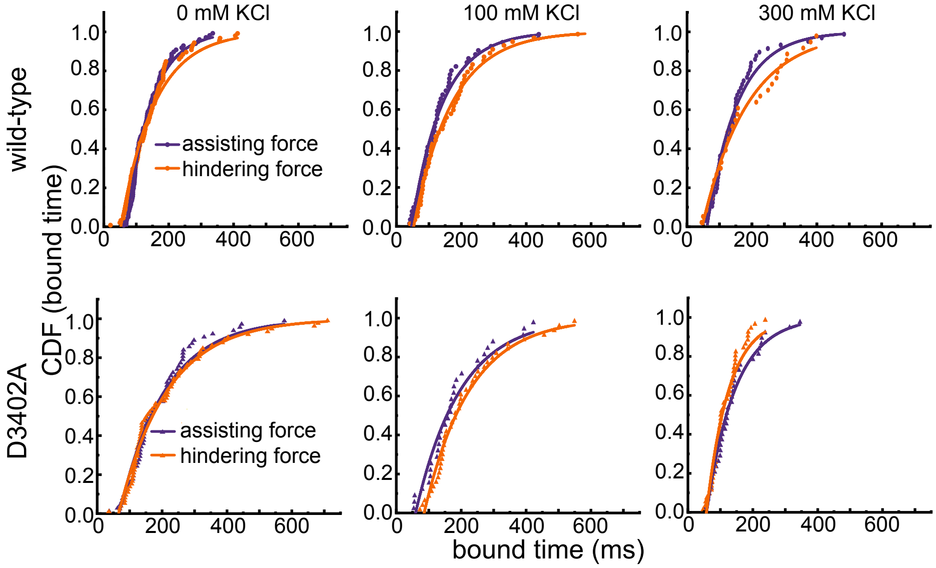


**Figure S5 Cumulative distributions of bound times for wildtype (*top*) and D3402A mutant (*bottom*) MTBDs on microtubules when subjected to increasing assisting (*purple*) and hindering (*orange*) loads with additional salt as indicated.** We performed K-S tests to test whether the assisting and hindering data were sampled from statistically different distributions, and we found p values > 0.05 in all cases. Fits to the assisting and hindering data (*purple* and *orange* lines, respectively) were done to Eq. 2 from the main text and are shown for clarity.

### Supporting Tables

**Table S2 Primers used for mutagenesis in this study**. Lowercase letters represent the altered bases making the mutations.

| Primer name | Sequence |
|---|---|
| MTBD D3402A Forward | TATGCAGcCATGTTAAAGCGAGTGGAGC |
| MTBD D3402A Reverse | ATTGAGCTGTGCAATCGCCCACTTCAC |
| MTBD D3320A Forward | GCTGGCTCTGgcgTCCATCTGCC |
| MTBD D3320A Reverse | TTCACAGCTGCAGGAGGG |

**Table S3 P-values of pair-wise two-tailed K-S test results for the distribution of bound times of the wild-type (WT) and D3402A mutant MTBDs on microtubules under load in the optical tweezer assay.**

| | | WT | | | D3402A | | |
|---|---|---|---|---|---|---|---|
| | [KCl] (mM) | 0 | 100 | 300 | 0 | 100 | 300 |
| WT | 0 | - | | | | | |
| | 100 | 0.32 | - | | | | |
| | 300 | 0.49 | 0.80 | - | | | |
| D3402A | 0 | < 0.001 | 0.0012 | 0.0096 | - | | |
| | 100 | < 0.001 | < 0.001 | 0.0080 | 0.77 | - | |
| | 300 | 0.33 | 0.14 | 0.077 | < 0.001 | < 0.001 | - |

Green shading corresponds to p values > 0.05 (not a significant difference)
Orange shading corresponds to p values < 0.05 (a significant difference)
